# Collective signalling drives rapid jumping between cell states

**DOI:** 10.1101/2023.05.03.539233

**Authors:** Elizabeth R. Westbrook, Tchern Lenn, Jonathan R. Chubb, Vlatka Antolović

## Abstract

Development can proceed in “fits and starts”, with rapid transitions between cell states involving concerted transcriptome-wide changes in gene expression. However, it is not clear how these transitions are regulated in complex cell populations, in which cells receive multiple inputs. It is also not clear to what extent these rapid transitions represent developmental commitment. Here we address these issues using *Dictyostelium* cells undergoing development in their physiological niche. A continuous single cell transcriptomics time series reveals a sharp “jump” in global gene expression marking functionally different cell states. By simultaneously live imaging the physiological dynamics of transcription and signalling over millimetre length scales, we show that the jump coincides with the onset of collective oscillations of cAMP, the positive feedback signal for multicellular development. Different jump genes respond to distinct dynamic features of signalling. The late gene expression changes of the jump are almost completely dependent on cAMP. In contrast, transcript changes at the onset of the jump require additional input. The spatial boundary marking the jump divides cells separated by only a few minutes of developmental time, with cells missing a jump then waiting several hours for the onset of the next wave of cAMP oscillations. This timing variability contrasts the prevailing developmental paradigm of a timed synchronous process and is associated with substantial pre-jump transcriptome variability. The coupling of collective signalling with gene expression is a potentially powerful strategy to drive robust cell state transitions in heterogeneous signalling environments. Based on the context of the jump, we also conclude that sharp gene expression transitions may not be sufficient for commitment.

## Introduction

The changes in gene expression occurring during developmental progression are not constant paced. In diverse developmental contexts, from plants, to *Dictyostelium,* to neurons, to adult and embryonic stem cells, developmental progression occurs by rapid and concerted transcriptome-wide switching from one gene expression state to the next (Antolovic et al., 2019; Artegiani et al., 2017; Giri et al., 2022; Moris et al., 2016; Nelms and Walbot, 2019; Rukhlenko et al., 2022; Saez et al., 2022). These rapid transitions imply a powerful and general mechanism for cells to robustly “commit” to a specific state in the presence of complex tissue signalling, by making cells insensitive to signals promoting alternative states, and by promoting coherence in the establishment of the new state.

Sharp switching between transcriptome states has usually been revealed by single cell transcriptomics methods. Although these approaches allow transcriptomes to be sampled from many cells at a time, and so enable classification of cell states, the measurements require disrupting the cells and their dynamic population structure. Consequently, it is unclear how rapid cell state switching is organised and coordinated in space and time within physiological cell contexts.

Here we investigate the coordination of rapid cell state transitions using the social amoeba, *Dictyostelium.* These cells enter their developmental programme upon exhaustion of their food source. After a few hours of starvation, cells begin signalling to each other using extracellular cAMP, which acts as a chemoattractant and drives the aggregation of the cells into a multicellular mound. Over the next 15-20 hours, the mound undergoes a series of morphogenetic transitions, resulting in the generation of the mature final structure-a fruiting body with spores suspended over the substrate by a stalk. In addition to these morphogenetic transitions, the cells change a substantial proportion of their transcriptome as they transition from the feeding state to the final structure. Time series analysis of transcriptomes at the population level reveals, as in other systems, that developmental progression is not constant paced (Parikh et al., 2010; Rosengarten et al., 2015). More recently, single cell transcriptome analysis of the mound stage revealed discrete states during the cell fate bifurcation process, indicating the concerted switching of the transcriptome within single cells (Antolovic et al., 2019).

Gene expression changes are regulated by a variety of signals: the onset of development is regulated by nutritional signalling (Jaiswal and Kimmel, 2019), quorum sensing (Clarke and Gomer, 1995) and cAMP (Cai et al., 2014; Corrigan and Chubb, 2014; Masaki et al., 2013), with other signals operating later during development (Williams, 2006). Despite the involvement of multiple signals during early development, most assays remove this signalling complexity, by plating cells from well-mixed cultures in non-nutrient buffer at uniform density. This removes the natural heterogeneity in developmental time within a *Dictyostelium* colony, and the complex external regulation experienced by each cell is reduced to a time-dependent wait for the onset of cAMP signalling.

To understand cell state switching in a more physiological context, we instead consider the early developmental programme in a mimic of the *Dictyostelium* physiological niche. The cells normally live in the soil, feeding on bacteria, and this is simulated in the lab by plating cells on a lawn of bacteria on an agar plate. As cells clear the bacteria, they create a plaque, in which the starving cells then undergo development. This niche-mimic contains the full asynchronous spectrum of developmental states, and more closely resembles the natural signalling complexity, in which nutrition (bacteria), cAMP and variations in cell density (quorum signalling) co-exist. We contextualise a sharp transition in transcriptome content-the jump-which occurs at the transition between the unicellular and multicellular stages of development. The jump emerges as a sharp spatial boundary in the colony as collective cAMP signalling begins. Jump gene expression requires cAMP signalling, however different jump genes respond to cAMP with different dynamic behaviours. Post-jump gene expression is almost completely dependent on cAMP, while early jump genes require additional signalling inputs. The jump differentially recruits cells separated by only minutes in developmental time, challenging the standard view of development as synchronous timer-based process. Based on the context of the jump, we infer that gene expression changes at the jump do not constitute commitment.

## Results

Understanding the regulation of cell state transitions during development requires defining cell states in the unperturbed physiological context. To define the major transitions during developmental progression, we collected a continuous single cell transcriptomics time course of *Dictyostelium* development. To capture development in a continuous manner, we collected cells from colonies of cells feeding on their bacterial food source (Figure 1A). In this context, cells feed on bacteria, and migrate further into the bacteria to acquire more food. Cells left behind starve, which triggers their developmental programme: single cells aggregate together by chemotaxis towards periodic signalling waves of cAMP, to form mounds. Subsequently, the mound goes through a series of morphogenetic steps, ultimately generating the final structure, with spores suspended above the substrate by a stalk. We collected a continuous streak of cells, from the bacterial zone through to the mounds, then generated single cell transcriptomes for 4743 cells.

**Figure 1.**
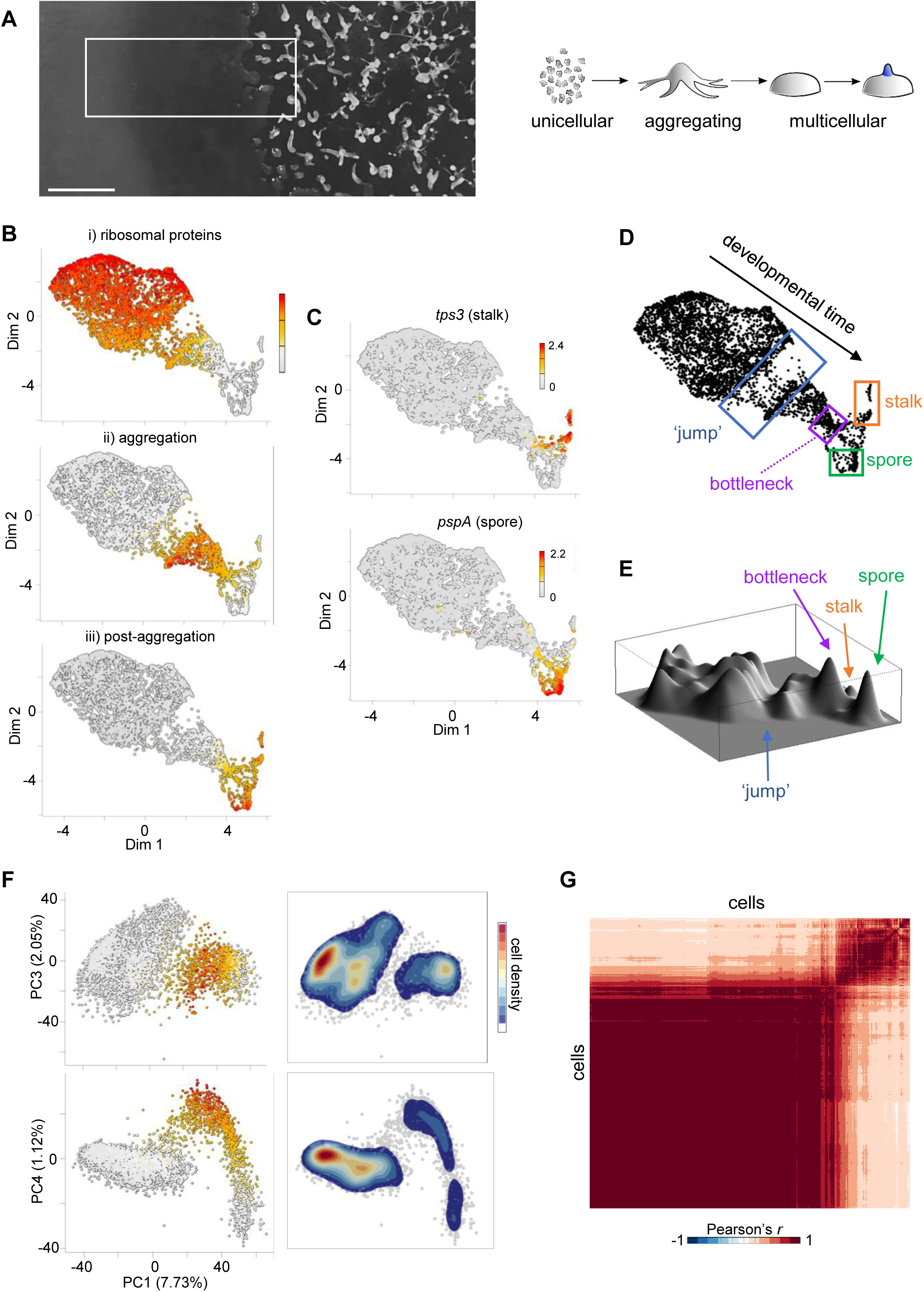
A jump in developmental progression. **A)** The *Dictyostelium* developmental niche. Left panel: cells are plated on a bacterial lawn with uncleared bacteria on the left. To the right, bacteria are cleared, cells enter the multicellular state, which goes through morphogenesis to the final fruiting body formation (far right). The white rectangle illustrates the continuous region sampled for transcriptome analysis. Right panel shows a schematic of the sampled life cycle stages. Scale bar 0.5cm. **B)** 4743 cells positioned in two-dimensional (2D) space, with each cell coloured by the mean expression of the following gene sets: i) ribosomal protein genes (78 genes), ii) aggregation genes (200) and iii) genes upregulated in aggregates (215). **C)** Expression of stalk (*tps3*) and spore (*pspA*) transcripts in 2D transcriptome space. Scale shows log10 of transcript counts (UMIs). **D)** Summary of transcriptome map, showing the jump, bottleneck and cell fate separation. **E)** Cell density landscape of D. Peak heights relate to cell abundance at specific transcriptome states. Few cells are found in the jump region, and cells accumulate in the bottleneck (Fig. S1D). **F)** Validation of the jump using PCA. Principal components (PCs) 3 and 4 are plotted against PC1. Each dot is a cell. Colours in left panels are the mean expression level of the aggregation gene set. Colours in the right panels correspond to relative cell density. PC1 approximates developmental progression. Separation of two cell populations (the jump) is clearly visible in both PC1-PC3 and PC1-PC4 space. Aggregation-specific gene expression increases just after the jump (see also Fig. S1E). **G)** Two main cell states revealed by a cell-cell correlation matrix, with two distinct clusters visible.

To visualise the data, we reduced its dimensionality to two components combining principal component analysis (PCA) and elastic embedding to retain both local and global data structure (Chen et al., 2019). To identify the direction of developmental time within the data, we labelled plots with panels of genes representative of specific stages of development (Figure 1B): the top plot shows expression of ribosome protein genes, which are strongly expressed during feeding but become repressed during starvation. The middle panel shows strong expression of aggregation-specific genes and the bottom panel displays expression of genes upregulated after aggregation. Expression of markers of the two principal fates, stalk and spore, occurs in the far right of the plot (Figure 1C). Overall, these data indicate developmental time proceeds from North West to South East along the backbone of the fish-shaped distribution (Figure 1D). This inferred directionality of developmental time is supported by overlaying expression of an independently generated population transcriptomic dataset (Katoh-Kurasawa et al., 2021) (Figure S1A and B), and expression of genes with cell cycle control functions, which label clusters in the undifferentiated zone and spore branch, consistent with known cell cycle activity (Muramoto and Chubb, 2008) (Figure S1C). The distributions of M and S-phase gene expression are broadly similar, consistent with studies showing *Dictyostelium* lack a G1 (Muramoto and Chubb, 2008; Zimmerman and Weijer, 1993).

### Cell state transitions during early development

The distribution of cells in this space reveals several key features. As cells differentiate, they encounter a region with few cells-“the jump”-indicating a rapid change in the global transcriptome of cells (Figure 1D). This “jump” is clearly observed in a 3D density plot of the data (Figure 1E), where peak heights correspond to cell density in gene expression space. After the jump, cells accumulate at a bottleneck, where their transcriptomes become similar, before undergoing a second rapid transcriptome remodelling, similar to the jump, as they separate into the spore and stalk fates, in agreement with earlier observations (Antolovic et al., 2019). Hierarchical clustering implies cells here proceed from the bottleneck into a mixed transcriptional intermediate state (Figure S1D; purple cells), in which spore and stalk markers can both be expressed, albeit with little overlap within individual cells, before the complete fate separation occurs.

In this study, we consider the first jump. To test if this jump is a biological effect or an effect of the two-dimensional data representation, we also represented the data using linear dimensionality reduction: PCA. In PCA, the PC1 axis reflects developmental time (Figure S1E), with the jump clearly apparent in several higher order principal components (Figure 1F), indicating it is not an artefact of the elastic embedding procedure. The jump is also clear in clustering of cell-cell Pearson correlations (Figure 1G), which reveals two major clusters, corresponding to the cells before and after the jump.

To gain insight into the gene expression changes occurring during the jump, we carried out unbiased hierarchical clustering on the whole dataset. The clustering revealed the sharp changes in global expression profiles occurring during the jump and identified four major clusters (Figure S1F left panel), which are highlighted on the 2D embedding plot (Figure S1F right panel): two clusters of cells before the jump and two after the jump. Based on gene expression signatures, these clusters represent cells that are feeding (red), starving (green), aggregating (blue) and mound stage (purple). Our inference here, also apparent in Figure S1A, is that the jump occurs at the onset of aggregation. This is consistent with population transcriptomic data based on morphologically-staged time series that show substantial transcriptome changes between single cell and multicellular stages (Katoh-Kurasawa et al., 2021; Parikh et al., 2010). However, the implication from our continuously-sampled data showing few cells caught within the jump is that the transition is concerted within individual cells, with two clearly demarcated attractor states (Figure 1D-F), features not resolvable using population average data. The majority of changes before the jump are repressive, with 66% of transcripts down-regulated in two waves before the jump (Figure S1F). Transcript clearance might result from transcriptional repression followed by constitutive RNA turnover, or by induced RNA decay. Concerted transcriptome shifts within a cell, based on transcriptional repression, would require the half-lives of repressed transcripts to be matched, to enable synchrony. This is not consistent with data showing a broad heterogeneity in turnover times for different mRNAs during starvation (Muramoto et al., 2012), implying the jump requires an induced RNA turnover mechanism.

To more precisely contextualise the jump with developmental progression, we used live imaging of transcription of jump marker genes, using transcriptional reporters inserted into endogenous gene loci. We identified jump markers in the transcriptome data that are representative of cells at different stages of the jump (Figure 2A). The *cafA* gene, which encodes a calcium binding protein, is induced prior to the jump. *carA,* the cAMP receptor gene, is expressed slightly later, with detectable induction before the jump. The *csbA* gene, which encodes a cell adhesion protein, is expressed post-jump. To directly visualise transcription of these markers during development, we inserted MS2 (Bertrand et al., 1998) and PP7 (Larson et al., 2011) stem loops into the endogenous gene loci, then used the cognate fluorescent MCP and PCP stem loop binding proteins to visualise nascent transcripts as spots at the site of transcription (Figure 2B) (Tunnacliffe and Chubb, 2020). Genes were imaged in pairs, with simultaneous imaging of both MS2 and PP7-tagged genes. To ensure physiological regulation, and to contextualise transcription with normal developmental progression, cells were directly imaged in the developmental colony. Both *cafA* and *carA* were strongly induced in cells before the onset of cell aggregation. In contrast, *csbA* only showed abundant transcriptional events in the zone of the colony undergoing aggregation.

**Figure 2.**
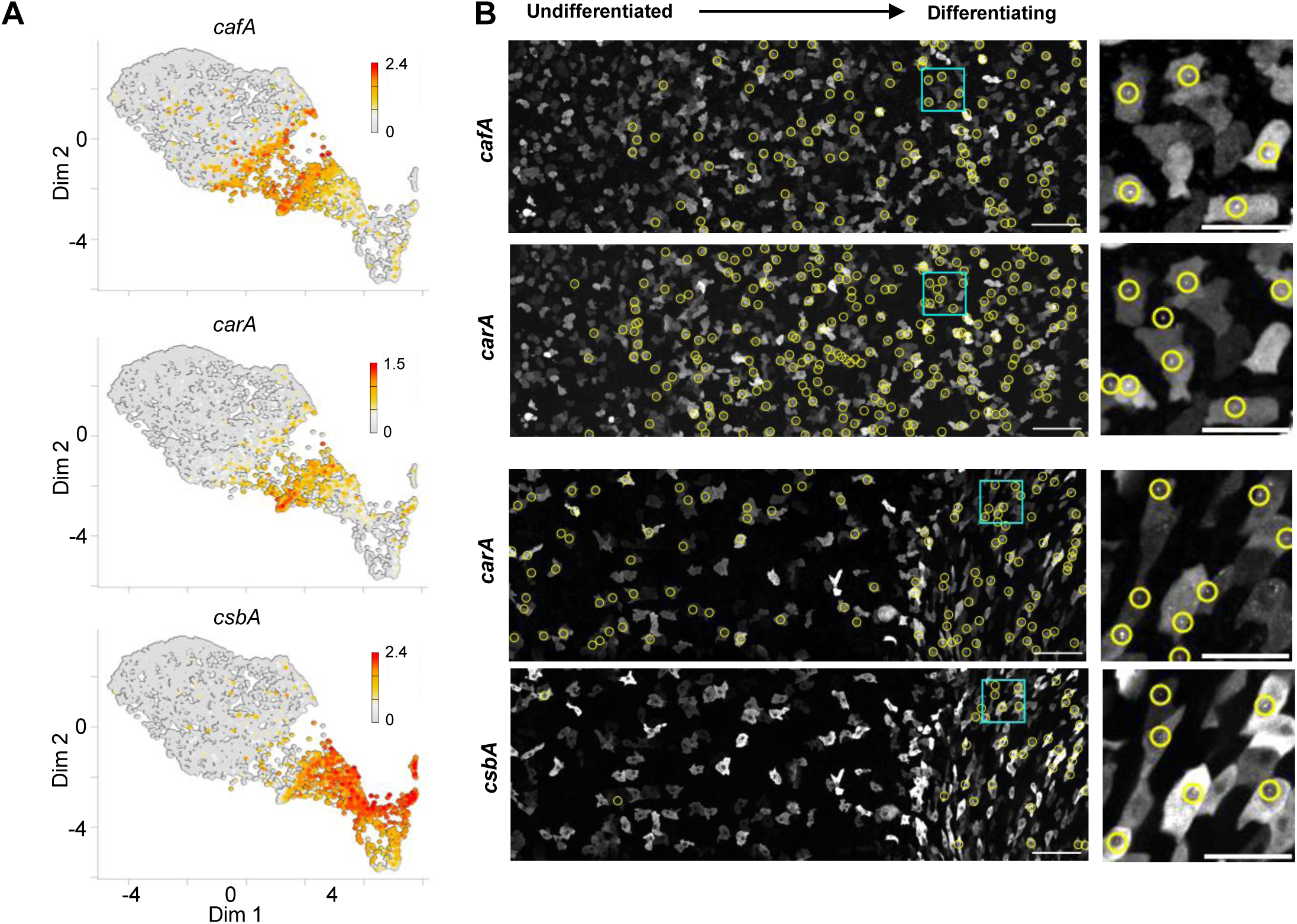
Matching jump transcripts to niche context. **A)** Expression of specific genes around the jump. 2D transcriptome maps are coloured by the mean expression level of the indicated genes. **B)** Imaging nascent transcription in the developmental niche of genes that change expression during the jump. On the left are the undifferentiated cells, on the right are cells beginning to show collective chemotaxis. Top two panels show transcription of *cafA*-PP7 and *carA*-MS2 in the same cells. Yellow rings highlight cells with spots corresponding to nascent transcription. Bottom two panels show transcription of *carA*-MS2 and *csbA*-PP7. Scale bars: 50 µm left and 20 µm right.

### Regulation of jump gene expression by cAMP

Developmental gene expression can be influenced by multiple signals, notably starvation time (Jaiswal and Kimmel, 2019) and extracellular cAMP (Cai et al., 2014; Corrigan and Chubb, 2014). To what extent are these signals, which are spatially heterogeneous in the niche, driving the gene expression changes at the jump? As both transcriptomics and the imaging imply jump genes such as *cafA* and *carA* are induced just prior or at the onset of aggregation, this suggested cAMP signalling may be responsible for the jump. To test this, we imaged transcription of jump genes in the colony (Movie S1), with parallel tracking of cAMP signalling, by mixing the transcriptional reporter cells with cells expressing the cAMP reporter, Flamindo2. Flamindo2 is an intensiometric cAMP reporter that dims in fluorescence when it binds to cAMP (Ford et al., 2023; Hashimura et al., 2019; Kundert et al., 2020)(Movie S2). We obtained simultaneous time series data recording the physiological dynamics of both transcription and signalling, in the unperturbed colony, from the undifferentiated cells through to the cells at the aggregation stage of differentiation (Figure 3A). Data are represented with the horizontal axis representing the position of each cell in the colony (Figure 3B). Undifferentiated cells are on the left, with the differentiating cells on the right. The vertical axis represents imaging time. For *carA*, the undifferentiated cells only showed sparse and sporadic transcription (Figures 3B and S2A), with transcription becoming strong and oscillatory in the more differentiated cells. The region of strong transcription coincided with the domain of cAMP fluctuations which showed oscillatory behaviour (Figure 3C-E, S2B-D), and continued as the signalling cells merged into an aggregate towards the end of the time series (observed as the constriction of fluorescence at the top right of Figure 3C). The *carA* expressing cells mark the zone in the colony where aggregating cells peel away from the rest of the population: the population that was spatially continuous at the onset of imaging separated gradually as the aggregate formed.

**Figure 3.**
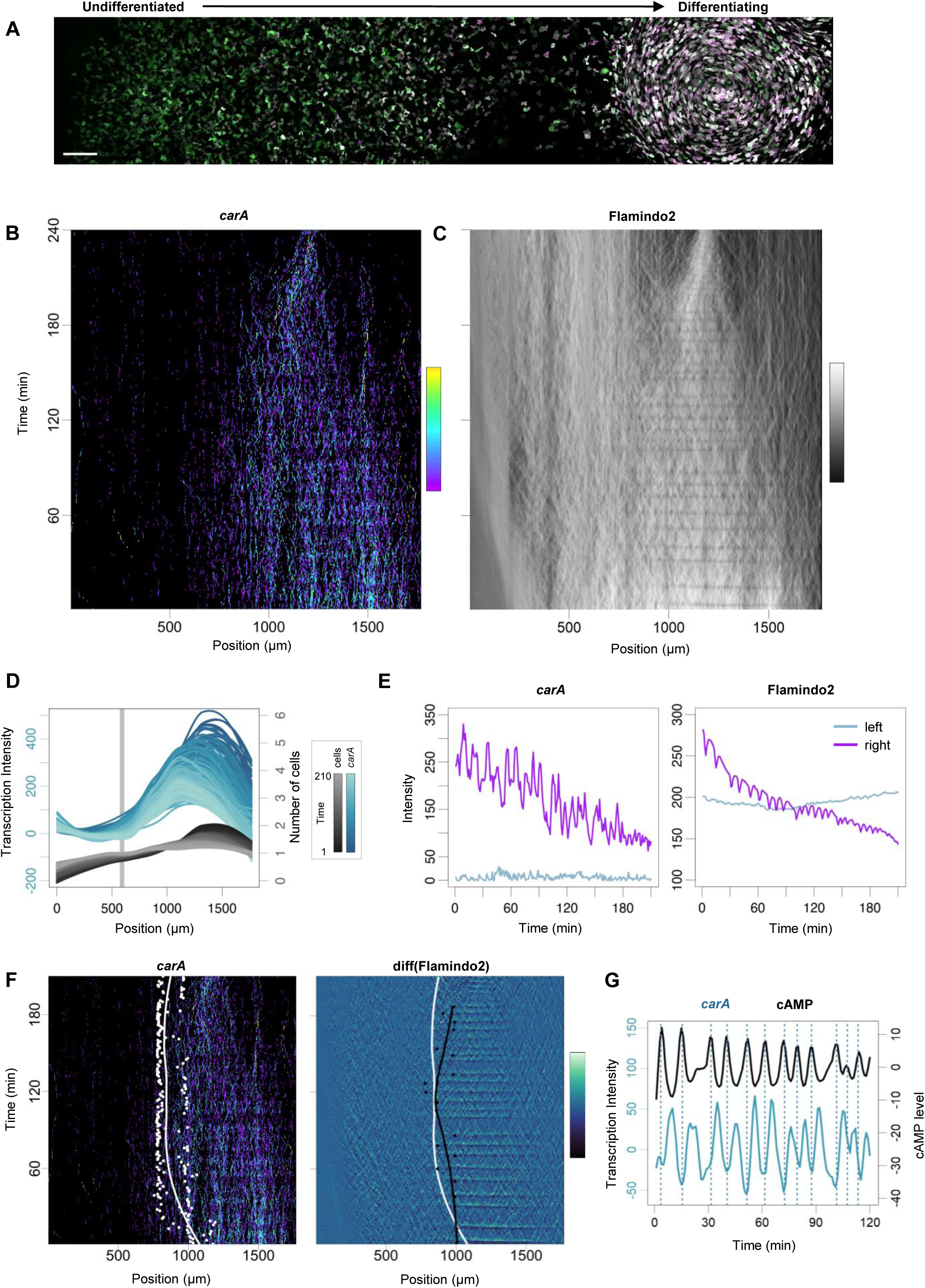
Coupling between jump transcription and signalling dynamics. **A)** Overview image of the *Dictyostelium* early developmental niche. Undifferentiated, feeding cells are on the left, becoming progressively more differentiated to the right entering a multicellular aggregate on the far right. Scale bar 100 µm. **B)** Imaging transcription dynamics of the jump gene, *carA*, in the niche. Horizontal axis reflects the axis of differentiation in A. The vertical axis is imaging time. Transcription spot intensity over time is shown, with activity level related by the colour scale bar. Transcription is sporadic in the less differentiated cells, becoming frequent and oscillatory as differentiation proceeds. Transcription spot intensities were averaged into 10 pixel bins (10 x 0.35µm). **C)** Same data as B, showing cAMP signalling using the Flamindo2 biosensor, which dims in fluorescence upon binding cAMP. Data show oscillations in differentiating cells. Cells merge into an aggregate towards the end of the time series. **D)** Increased transcription activity during differentiation. Plots summarise data in B, and also shows the distribution of cells in the population. Changing transcription and cell distributions over time are shown as different colour shades (see colour scale). The grey line corresponds to the minimum in cell density, where the population splits during the transition to multicellularity. **E)** Transitions in transcription and signalling dynamics across the niche. Left panel shows distinct *carA* transcription dynamics comparing zones left and right of the grey line in D. Right panel shows the distinction between oscillatory and non-oscillatory cAMP dynamics either side of the grey line. **F)** Positional coupling between transcription and signalling dynamics. Left-white spots areinflections of the curves of transcriptional intensity values at each imaging time point. The white line is a regression line summarising the distribution of points. Right-black dots show inflection points for cAMP signalling, with the black line the regression line and white line the same as in the left panel. Inflection values were calculated at timepoints of cAMP wave maxima. Diff(Flamindo2) represents the difference in intensity between one time point and the subsequent one. **G)** Temporal coupling between transcription and signalling oscillations. Peaks in cAMP signalling (vertical lines) occur 4-5 minutes prior to peaks in *carA* transcription.

To what extent are the oscillations temporally and spatially coupled? A quantitative analysis of *carA* transcription indicated *carA* induction occurs at the same region at which cAMP relay is occurring. This was revealed by substantial overlap in the inflections of the curves summarising transcription and signalling activity (Figure 3F, S2E). The period of the cAMP and transcription oscillations was similar, however the phases of transcription and signalling waves are offset (Figure 3G, S2F). A cross correlation analysis revealed a lag of 4-5 minutes between the peak of the cAMP wave and the peak of transcription, possibly reflecting signalling lags from receptor to gene, such as transcription factor shuttling times (Cai et al., 2014), in addition to the time for transcripts to build up at the locus. Overall, these data imply induction of *carA* by collective cAMP signalling.

To directly test the role of cAMP in inducing jump transcription, we imaged *carA* transcription together with signalling in *acaA-* mutants (Figure 4A), which have lost the adenylyl cyclase that synthesizes cAMP during early development. These mutants show a loss of cAMP signalling using the Flamindo2 reporter (Figure 4A right panel). The rare sporadic *carA* transcriptional events were still observed, with slightly enhanced activity further from the bacterial zone, however the gene did not show the strong induction of transcription observed in the wild-type developmental collective.

**Figure 4.**
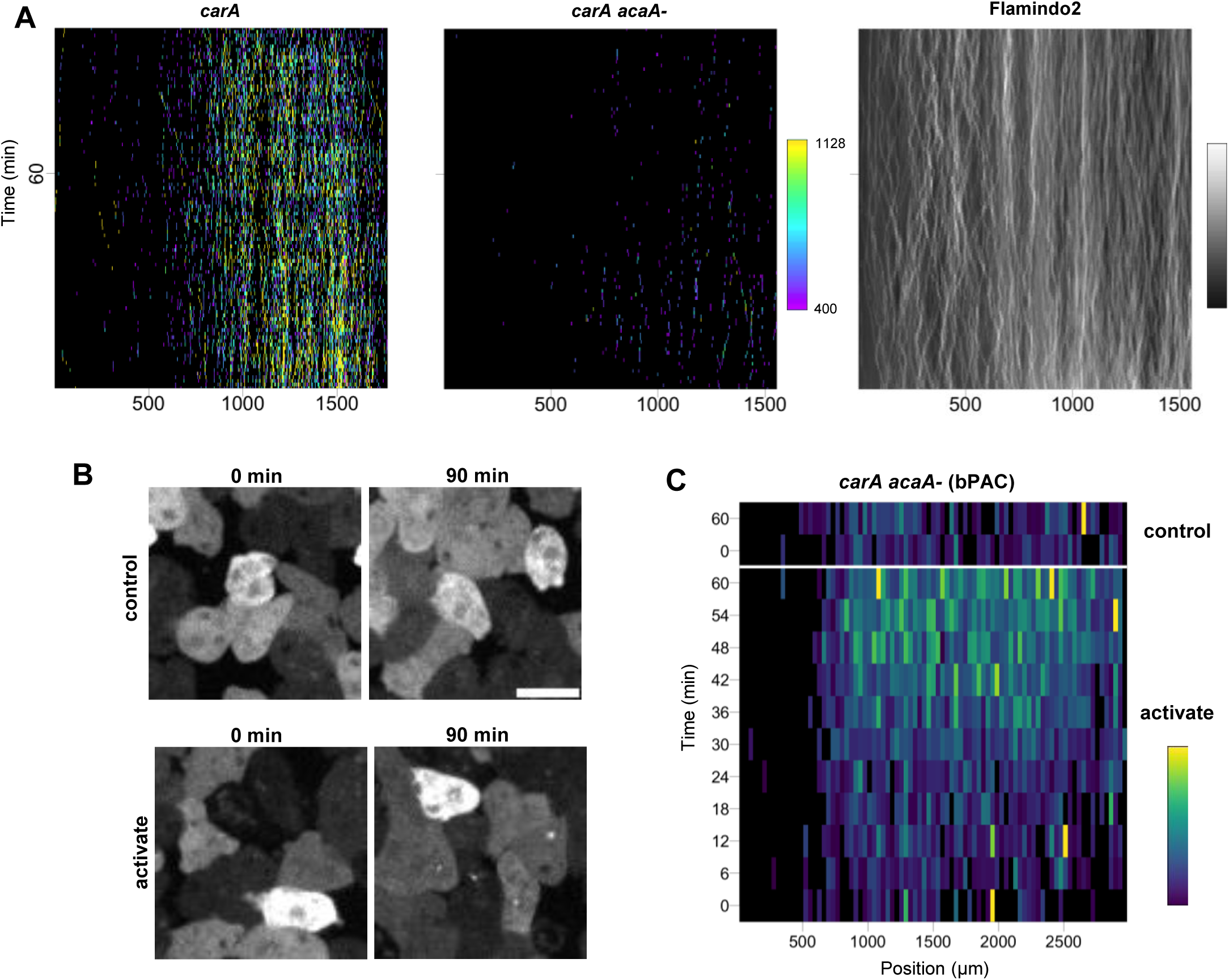
Functional coupling of cAMP signalling and jump gene expression. **A)** Loss of *carA* transcription in cells lacking a functional adenlyl cyclase A (ACA) gene. Left panel shows the rapid increase in *carA* transcription in wild-type cells (expanded view of Figure 2B). Central panel shows loss of *carA* induction in *acaA-* cells. Right panel shows absence of cAMP oscillations in *acaA-* cells. Typical experiments shown. 3 wild-type, 3 *acaA-* biological repeats carried out. **B)** Optogenetic rescue of jump gene expression: *acaA-carA-*PP7 cells mixed with *acaA-* cells expressing optogenetic adenylyl cyclase, bPAC. Cells were pulsed with blue light at 6 minute intervals to mimic normal cAMP signalling. Strong induction of transcription was observed in pulsed cells (bottom) compared to non-pulsed cells (top). Scale bar 10 µm. **C)** *carA* shows context and time-dependent responses to exogenous induction of cAMP using bPAC. Heatmap shows *carA* induction in the cell population after 30 minutes of pulsing, but not close to the undifferentiated zone. Transcription spot intensities were averaged into 100 pixel bins (100 x 0.35µm). Typical experiment is shown from 3 repeats (2 biological).

To test to what extent cAMP signalling is sufficient to induce jump gene expression, we exposed cells across the colony to periodic pulses of cAMP using the optogenetic adenylyl cyclase, bPAC from the soil bacterium *Beggiatoa* (Stierl et al., 2011). To effectively control the experiment in the absence of exogenous cAMP signalling, we used *acaA-* cells, to prevent cAMP signals propagating across the colony, and to allow test (activated) and control (not-activated) cells to be compared in the same conditions. We activated bPAC at 6 minute intervals with blue light pulses along the entire starvation axis of the developmental collective. This pulse frequency was used to mimic normal excitable cAMP signalling pulses. This regime of pulsing caused the induction of bright *carA* transcription spots in activated cells, but not in no light controls (Figure 4B). When examined over the whole collective, the induction process revealed other features of the niche that influence cell responsiveness (Figure 4C). Firstly, the induction was not immediate-the cells required around 30 minutes of pulsing before showing strong induction, indicating some requirement for priming. Secondly, the induction was spatially restricted, with the less differentiated cells at the left of the colony not showing *carA* induction, implying repression by some feature of cell context in this zone. So although these data indicate oscillatory cAMP signalling drives the jump, the responsiveness of cells depends on their context in the niche.

### Jump genes have different regulatory inputs

Temporal coupling was observed between cAMP oscillations and the *cafA* gene (Figure 5 and S3), however *cafA* showed different behaviour compared to *carA*. Transcription of *cafA* was observed in areas of the cell population without oscillatory cAMP signalling, although stronger transcription was observed in the zone where cAMP oscillations were detected (Figure 5A-C). The transcription was also oscillatory, however, unlike *carA,* the gene was repressed at the higher cAMP oscillation frequencies occurring later in the time series. A further difference between *carA* and *cafA* was apparent in the offset between transcription and signalling, with *cafA* transcription maxima delayed from cAMP maxima by 90 seconds or less (Figure 5D, S3F).

**Figure 5.**
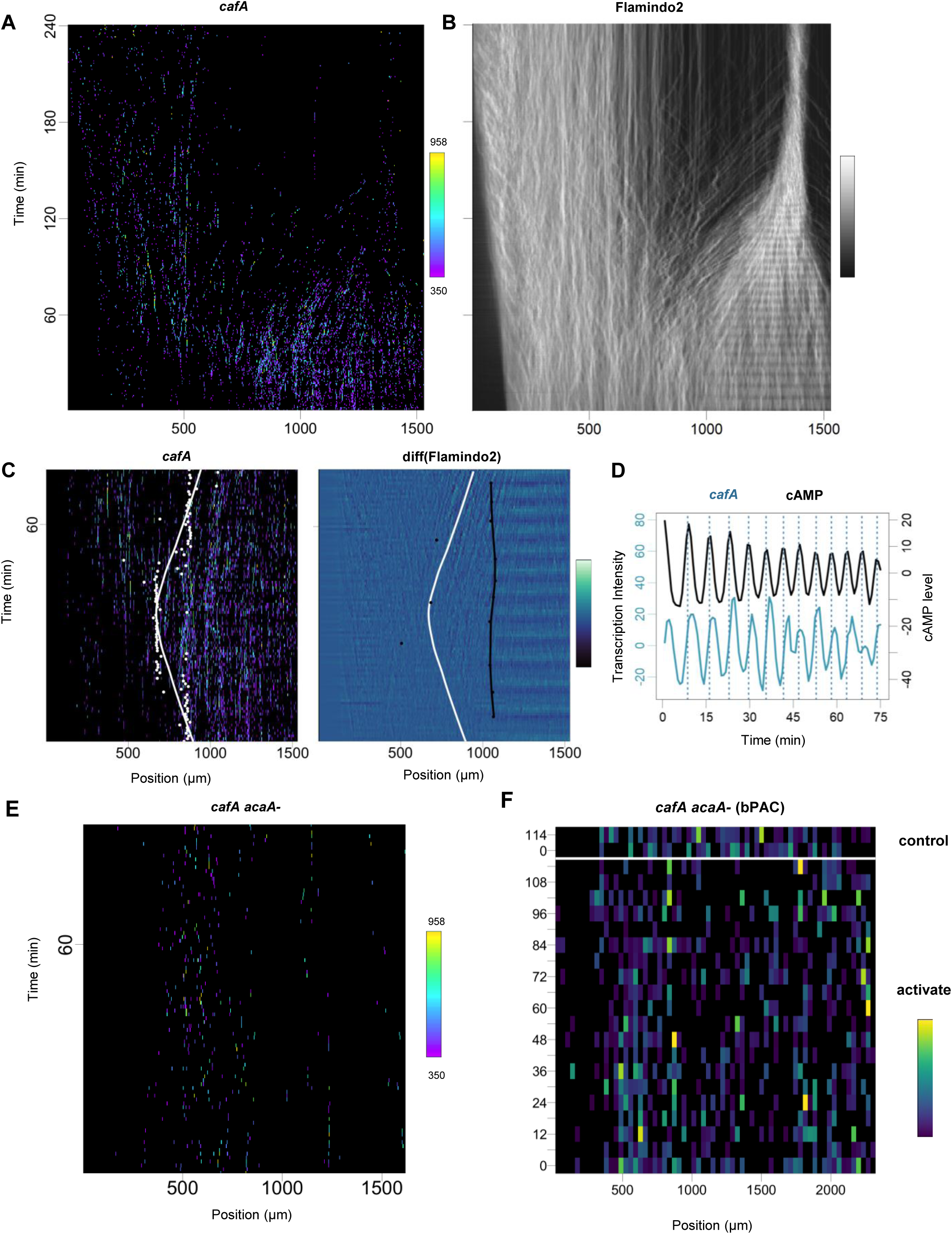
Alternative coupling strategies between jump signalling and transcription. **A)** Imaging transcription dynamics of the jump gene, *cafA*. Horizontal axis reflects the axis of differentiation. Vertical axis shows imaging time. Transcription is sporadic in the less differentiated cells, becoming frequent and oscillatory during differentiation. **B)** Same cell field as A, showing cAMP signalling dynamics. Cells merge into an aggregate during the movie. **C)** Non-overlapping boundaries of transcription and signalling. Left-white spots represent inflections of the curves of transcriptional intensity values at each imaging time point. The white line is a regression line summarising the distribution of points. Right-black dots show inflection points for cAMP, black line is the regression for these points. White line the same as in left panel. **D)** Temporal coupling between transcription and signalling. Peaks in cAMP signalling slightly precede peaks in *cafA* transcription. **E)** Loss of *cafA* transcription in cells lacking a functional adenlyl cyclase A (ACA) gene. Typical experiments are shown in A and E from 7 wild-type and 4 *acaA-* (biological repeats). **F)** Optogenetic activation of cAMP does not rescue *cafA* gene expression: *acaA-cafA-*PP7 cells mixed with *acaA-* cells expressing optogenetic adenylyl cyclase, bPAC. Cells were pulsed with blue light at 6 minute intervals to mimic normal cAMP signalling. Unlike for *carA,* induction of transcription was not observed in pulsed cells. Transcription spot intensities were averaged into 100 pixel bins (100 x 0.35µm). Typical experiment is shown from 7 repeats (3 biological).

To directly test the role of cAMP in inducing *cafA* transcription, we imaged *cafA* transcription and cAMP signalling in *acaA-* mutants (Figure 5E). The rare sporadic *cafA* transcriptional events were still observed. However, the gene failed to show the strong induction of transcription normally observed in wild-type cells. Unlike *carA,* the *cafA* gene was not induced by pulsed optogenetic synthesis of cAMP (Figure 5F). Therefore, although strong induction of both genes requires cAMP, *carA* and *cafA* show distinct kinetics of coupling to cAMP signalling.

The coupling of *cafA* transcription to cAMP may follow the rules inferred for the transcriptional oscillations of the *csaA* gene (Cai et al., 2014; Corrigan and Chubb, 2014). With the caveat that *csaA* oscillations were observed with cells differentiating in buffer, rather than in the niche, the gene was proposed to show two step regulation, with activation and repression at different stages of the cAMP oscillation cycle. The effect of this scenario is that the gene is switched off at high cAMP wave frequencies, as the repression occurs before the activated state has sufficient time to be productive. Transcription of *cafA* is repressed at high cAMP frequencies (Figure 5), in addition to showing activation independent of cAMP oscillations, much like *csaA.* In contrast, the *carA* gene is not inactivated at high signal frequencies (Figure 3), suggesting a more simple one step model, in which the gene activation mirrors the level of cAMP (with a lag) but no explicit repressive input.

We then further tested the requirement for cAMP signalling for a broad set of genes changing their expression at the jump. To define this set of genes, we intersected high temporal resolution population transcriptomic datasets from synchronous developmental protocols (Katoh-Kurasawa et al., 2021) with our own continuous single cell transcriptome data from the physiological colony (Figure S4). We categorised genes into three profiles: repressed at the jump (pre-jump genes), induced spanning the jump (“jump” genes such as *carA* and *cafA*) and induced after the jump (post-jump). Comparing with population transcriptomic data on wild-type and *acaA-* cells reveals effects of cAMP removal on all three categories. 85% of pre-jump genes fail to be repressed without cAMP signalling (Figure S4B; 46/54 genes). For jump genes, 16/22 showed partially reduced expression, with the remainder losing induction completely (Figure S4C). Post-jump genes almost entirely showed complete lack of normal developmental expression, with only 1/82 genes (*csbC)* retaining detectable induction (Figure S4D). Overall, these data indicate that erasure of the undifferentiated state requires cAMP and the post-jump state is effectively absent without cAMP. In contrast, as also indicated by our niche-based imaging approaches, induction of the transcripts spanning the jump state requires a mixture of cAMP signalling and other inputs.

### Collective signalling separates cells of a similar developmental age

In the developmental niche, cells that peel off to join streams of aggregating cells are initially spatially directly adjacent to cells of a similar developmental time (eg. Figure 3B). To quantify this, we captured low magnification time series of the developmental colony (Figure 6A). The cells advance into the bacterial zone at a constant rate of around 1.9 µm min^-1^, which is slightly slower than they migrate *in vitro* in buffer (Chubb et al., 2002). The events in which cells peel off to form streams and then mounds occur around once every 4 hours (Figure 6B,C) although this can be as much as 10 hours. This may be an underestimate of the variability, with rare mounds forming well behind the normal band of mound formation, in the zone containing fruiting bodies. Overall, this variation implies the absolute time of starvation, which reflects the continuous clearance of the bacteria away from the starving cells, is not a precise predictor of the time at which the cell enters multicellular development, which is a discrete event. As a consequence, cells entering mounds will vary in developmental time by the size of the interval between peel-off events. To contextualise this variation in timing-the normal starvation time before aggregation onset in synchronous developmental protocols is 4-6 hours, depending on the strain used, and normal experimental variation. As a result, cells entering late into a mound will have experienced around two fold (or sometimes considerably more) extra nutrient deprivation than cells early into a mound (Figure 6D). This represents a substantial spontaneous heterogeneity in cell signalling history, which may underlie the observed sub-clustering of cell gene expression states just before the jump (Figure 1E). This heterogeneity may have functional consequences: nutrient-deprived cells tend to adopt the stalk rather than spore fate (Thompson and Kay, 2000). The spontaneous formation of mixed-age mounds by the jump would therefore provide a straightforward source of nutritional heterogeneity to facilitate robust cell type patterning.

**Figure 6.**
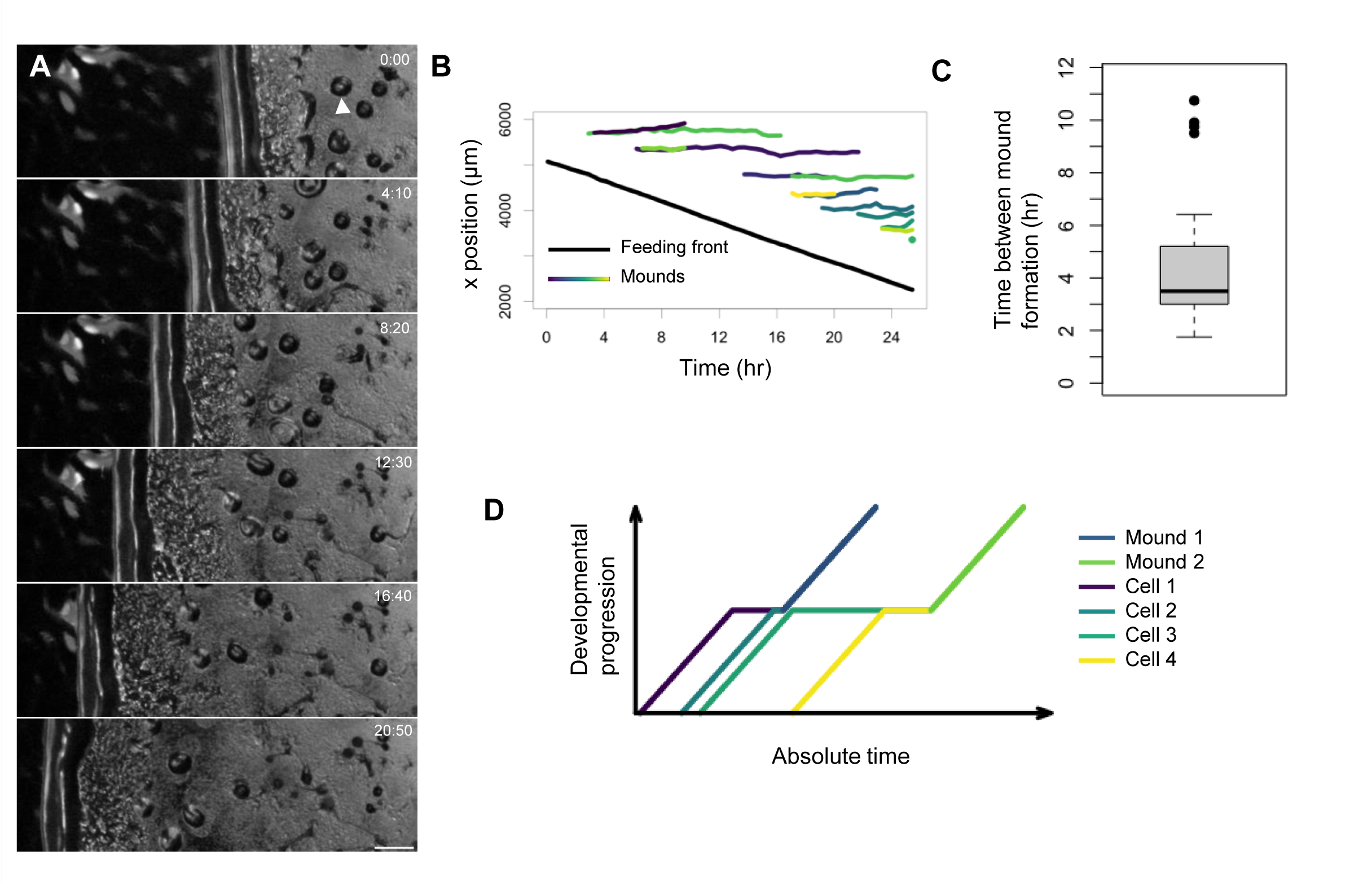
Aggregation combines cells with widely different developmental times. **A)** Cell deposition by the advancing feeding front. Stills from a time-lapse movie show cells clear the bacterial lawn at a constant rate from right to left. Behind the cleared lawn cells are left behind, which aggregate into mounds (arrowhead) at discrete steps. Scale bar 0.5 mm. **B)** Quantification of the movie in A showing feeding front progression (black line) and individual mound formation events in the recently starved zone. For clarity, the data are extracted from the bottom half of each panel in the movie. Different mounds shown as different colours. **C)** Time between mound formation within 150 pixel strips (0.63 mm). **D)** Scenarios caused by discrete budding events. Cells are continuously shed from the feeding front. Each line represents an example cell that leaves the feeding front from this continuously shed population. The purple cell leaves the front and forms a mound with the blue cell. The green cell buds shortly after the blue cell but waits a long period to enter a mound with the yellow cell.

## Discussion

There are key features of cAMP signalling that well suit its ability to drive a sharp change in cell state. As with many tissue signalling processes in more complex systems (Deneke and Di Talia, 2018; Dieterich et al., 2006; Ender et al., 2022; Liu et al., 2022; Pond et al., 2022), signalling by cAMP is excitable: as one cell is activated, it releases more signal to its neighbours, which then further spread the signal (Ford et al., 2023; Gregor et al., 2010; Tomchik and Devreotes, 1981). This signal relay will enable coordinated switching of a cell population into the new state, necessary for an organised response. In addition, the genes induced at the jump, as exemplified by *carA* (which encodes the cAMP receptor) provide the potential for positive feedback. The ability of a signal to induce its own receptor, in addition to the induction at the jump of other genes required for cell aggregation, will further strengthen cAMP signalling between cells. This mutual interaction allows an amplification ideally suited to rapidly lifting a cell out of one state and into the next.

One consideration is that cells would need to be able to perceive cAMP to get the amplification process started, which will require a cAMP receptor. Consistent with this requirement, feeding cells can show basal levels of expression from the *carA* locus (Muramoto et al., 2012), so there will be the potential to detect early arriving cAMP. A further issue is that although induction of transcript clearance and post-jump transcription appear dominated by cAMP regulation, the induction of most genes spanning the jump, notably *cafA*, is also modulated by other inputs. This makes regulatory sense-for a cell to embark on a sharp state transition, multiple inputs would provide more robustness to this decision. Overlying a collective signal over a timing mechanism (starvation) means the cell will only jump when there is a sufficient quorum to make the transition to multicellularity worthwhile, whilst allowing sufficient time to not miss out on another opportunity to feed.

Sharp state transitions or jumps have been implicated as “commitment” points (Mulas et al., 2021). Definitions of commitment vary, but a standard usage implies some resistance against cells reverting to their former state. This usage may not apply to the jump we are considering here. Differentiating *Dictyostelium* cells can de-differentiate rapidly in response to the reapplication of their nutrition source (Finney et al., 1987; Nichols et al., 2020). De-differentiation of most cells in the population is complete within no more than a day and this applies to cells up to the point of terminal differentiation-many hours after the jump. This indicates the jump itself presents no absolute barrier to cell state reversion. However, de-differentiation is usually induced by experimental disaggregation of developing structures. If nutrition is applied to intact structures, or cells around the onset of multicellularity, then they de-differentiate poorly, if at all (Katoh et al., 2007). This resistance to dedifferentiation can be considered commitment but is likely to result from the stability of the signalling across networks of cells, rather than any stable cell autonomous state resulting from the jump. Indeed, mutant cells which generate unstable mounds show signatures of de-differentiation (Katoh-Kurasawa et al., 2021), suggesting the differentiating state is stabilised by cell interactions, not directly by gene expression state. Does this relate to cell state transitions in general? To an extent, perhaps. Ground state mouse embryonic stem cells can populate preimplantation blastocysts with high efficiency-yet slightly more differentiated cells can reset with a low frequency to contribute to chimeras, although most are lost by cell competition (Alexandrova et al., 2016). Another consideration is that development often requires much more time, and perhaps more cell state transitions than *Dictyostelium* development, so developmentally advanced cells may no longer have the machinery to interpret the signals promoting an earlier state, meaning de-differentiation can only be enforced by more aggressive approaches, such as forced transcription factor expression.

The single cell gene expression data reported here reveal unexpected sources of cellular heterogeneity. Whilst the effects of dimensionality reduction need also to be considered, the cells appear heterogeneous in the feeding state before becoming more heterogeneous prior to the jump. This increase in heterogeneity was previously observed in cells differentiating in buffer, which was suggested by modelling to result from the effect of transcription repression on transcriptional noise (Antolovic et al., 2017). Starving cells reduce their overall transcriptional output (Mangiarotti et al., 1981), as might be expected in a context opposed to extensive biosynthesis, which may provide the driver for the increased noise. We show here that there is another potential layer of heterogeneity arising from differences in starvation time of cells undergoing the jump. Based on the effects of experimental nutrition deprivation on perturbing cell fate allocation (Thompson and Kay, 2000), this spontaneous heterogeneity in nutritional history for cells entering the multicellular stage may contribute to the overall fate diversity between cells in the final developed structure. Input to fate choice will also likely include differences in cell cycle position, which can be a functional source of heterogeneity for cell type allocation in *Dictyostelium* (Gomer and Firtel, 1987; Thompson and Kay, 2000) and other differentiation systems (Pauklin and Vallier, 2013).

While the pre-jump heterogeneity is largely consistent with the long-held notion that fate choice during development requires differences between cells in feeding and starvation, it is not clear why this heterogeneity should then become reduced before the onset of fate marker expression-the bottleneck. This constriction of cell variability resembles previous single cell transcriptome measurements in the mound (where cell fate bifurcation first becomes detectable), which identified a compact population before the branching into spore and stalk trajectories (Antolovic et al., 2019). The most likely explanation is that the cells at this stage are aggregating or recently aggregated and regardless of their final fate, will be challenged with expressing the components required for enacting the single cell to multicellular transition. These transcripts dominate the measured transcriptome and are shared by all cells (Antolovic et al., 2019). Based on the sensitivity of single cell transcriptomics, these might be expected to obscure the more variable transcripts conveying information to cells for fate allocation. Alternatively, the cell-cell differences required to inform fate may be better represented in the proteome.

## Materials and Methods

### Cell handling

Cells were cultured on lawns of *Klebsiella pneumoniae* on plates of SM agar (Urushihara, 2006). For transcriptomics, we used the wild isolate strain NC4. For genetic modification, we used the Ax3 strain, and Ax3 expressing the nuclear marker, H2Bv3-mCherry under the control of the endogenous *rps30* promoter (Corrigan and Chubb, 2014).

### Molecular biology

To image transcription, PP7 repeats were inserted into endogenous *cafA, carA* and *csbA* genes. Fragments containing 24 PP7 repeats and a blasticidin resistance gene were inserted between promoter and coding sequences. Dual transcriptional reporter cell lines with *carA*-MS2 and either *cafA-*PP7 or *csbA-PP7* were generated in Ax3 *carA*-MS2 knock-in cells (Muramoto et al., 2012). Single reporter lines for *carA-*PP7 and *cafA-*PP7 were generated in H2Bv3-mCherry labelled cells. To detect MS2 and PP7 stem loops we expressed GFP and TdTomato-tagged MCP and PCP proteins (Antolovic et al., 2019). For stable uniform Flamindo2 expression, we targeted a codon-optimised Flamindo2 gene into the *act5* gene of Ax3 cells as previously described (Ford et al., 2023). To disrupt the *acaA* gene, we used the *acaA* targeting vector from (Tweedy et al., 2020). For bPAC, a codon-optimised bPAC gene (Ford et al., 2023) was expressed from the extrachromosomal vector pDM1203 (Paschke et al., 2018), in Flamindo2-expressing *acaA-* cells.

### Single cell transcriptomics

For a continuous scRNAseq timecourse, we took a scrape of feeding fronts of NC4 cells, from inside the bacterial zone through to the mound stage of development. Cells were inoculated into ice-cold KK2 buffer (20mM KPO_4_, pH 6.0), and disaggregated by gentle pipetting. To remove bacteria, cells were centrifuged at 720g for 2 minutes, then resuspended in ice-cold KK2. Single cell transcriptomes were derived using the Chromium Single Cell A Chip platform (PN-1000009) based on (Nichols et al., 2020). Detailed information on sequencing, downstream processing and data analysis is in the Supplementary Methods. Transcriptomes from 2671 and 2072 cells, from two replicates, were used for further analysis. Sequencing data are available at GEO: GSE220242. Code for scRNAseq data analysis is available at https://github.com/Vlatka22/scRNAseq_Pipeline.

### Imaging and image analysis

For imaging gene activity with signalling, transcriptional and signalling reporter cells were mixed at a 1:2 ratio, and spotted onto lawns of *Klebsiella* on diluted SM agar plates (1 SM : 19 H_2_O). After 3 days for colonies to form, agar pads were excised and inverted onto imaging dishes (µ-Dish, Ibidi, 81156). Imaging used an inverted spinning disk confocal microscope (3i) using a 63 x oil lens, with a Prime 95B CMOS camera (Photometrics). We captured 14 to 16 z slices, with a 0.4 µm step size and 2x2 binning. 3D stacks were captured every 45 or 60 seconds at multiple xy positions across the cell population, with fields of view stitched to generate a complete view of the early developmental niche. GFP and mCherry/TdTomato were excited with 488 nm and 561 nm lasers, respectively, with laser powers optimised for best resolution alongside maintained cell health. For bPAC activation, transcriptional reporter cells were mixed 1:2 with bPAC-expressing cells. Activating bPAC used a 3D stack with a 488 nm laser every 6 minutes. This illumination had the dual function of activating bPAC and collecting transcription spot data.

For low magnification imaging of feeding front dynamics and mound formation, we spotted AX3 cells on bacterial lawns on 1:5 diluted SM agar plates, allowed 3 days for colonies to form then captured images every 5 minutes for up to 25 hours. Images were captured using a Dino-Lite digital microscope version 2.0 in a humidified chamber. We tracked the x position of the feeding front and mound position every 5 frames.

Spot detection was based on the approach from (Corrigan et al., 2016). To identify cAMP waves, we masked signal from the transcriptional reporter cells, which are more variable in their background intensity than the Flamindo2 cells. The intensity of the remaining cell-containing pixels (representing primarily the Flamindo2 signal) was averaged at each timepoint. Detailed analysis protocols and methods to compare signalling and transcription distributions are described in the Supplementary Methods. Image analysis code containing links to spot and signal intensity data is accessible at https://github.com/Vlatka22/ImageData_Analysis.

## Acknowledgements

This study was supported by a Wellcome Trust Senior Fellowship (202867/Z/16/Z) to JRC and a PhD studentship to ERW from MRC funding to the LMCB (MC_U12266B). The funders had no role in study design, data collection and interpretation, or the decision to submit the work for publication.

## Author Contributions

Conceptualization, ERW, TL, JRC, VA.; Methodology, ERW, TL, JRC, VA.; Investigation, ERW, TL, JRC, VA.; Writing – Original Draft, JRC, VA, ERW; Writing – Review & Editing, JRC, VA, ERW, TL; Funding Acquisition, JRC.; Software, VA, ERW, TL.; Formal analysis, VA, ERW; Supervision, JRC, VA.

## Declaration of Interests

The authors declare no competing interests.

## Supplementary Methods

### Cell handling

Cells of the wild isolate strain NC4 were cultured on lawns of *Klebsiella pneumoniae* on plates of SM agar. For genetic modifications, we used the Ax3 strain, and an Ax3 strain previously engineered to express the nuclear marker, H2Bv3-mCherry, under the control of the endogenous *rps30* promoter (Corrigan and Chubb, 2014). For DNA transformations, we used electroporation protocol based on H50 buffer (Paschke et al., 2018), with selections in standard HL5 axenic growth medium at 22°C, in tissue culture dishes. Selection used 20 µg/mL G418 for extrachromosomal expression vectors and either 10 µg/mL blasticidinS or 35 µg/mL hygromycin for gene targeting vectors.

### Molecular biology

For targeting with PP7 cassettes, fragments containing 24 PP7 repeats (Larson et al., 2011) and a blasticidin resistance (bsr) gene (Faix et al., 2004) were inserted between promoter and coding sequences of *cafA, carA* and *csbA* genes. For *carA-*PP7, we used the *carA-*MS2 targeting vector described previously (Muramoto et al., 2012), and replaced the BamHI fragment containing MS2-bsr with a BamHI fragment containing PP7-bsr. For *cafA-PP7*, we generated a targeting vector with targeting arms cloned as follows: -297 to +281 (promoter, with +1 marking the ATG), +284 to +1310 (coding sequence); for *csbA*: -373 to +274 and +288 to +1148, with HindIII and BsrGI used for cloning promoters, and SpeI and NotI for coding sequences, with PP7-bsr inserted using BsrGI and SpeI. Dual transcriptional reporter cell lines with *carA*-MS2 and either *cafA-*PP7 or *csbA-PP7* were generated in the Ax3 *carA*-MS2 knock-in cell line, pre-modified by Cre recombinase expression to remove the bsr. Single reporter lines for *carA-*PP7 and *cafA-*PP7 were generated in the H2Bv3-mCherry labelled Ax3 cells. Correct single copy insertions were validated by Southern blotting. Labelling of the MS2 and PP7 repeats was enabled by expression of extrachromosomal plasmids expressing GFP or TdTomato tagged MCP and PCP stem loop binding proteins. For stable even expression of Flamindo2, we targeted a codon-optimised Flamindo2 gene into the *act5* gene of Ax3 cells as previously described (Ford et al., 2022). To disrupt the *acaA* gene, we used a hygromycin-based *acaA* targeting vector in the transcriptional reporter cells, Ax3 cells and Flamindo2 AX3 cells. For bPAC, a codon optimised bPAC gene was expressed from the extrachromosomal vector pDM1203(Paschke et al., 2018), in Flamindo2 expressing cells with a disrupted *acaA* gene.

### Single cell transcriptomics

For single cell RNAseq of a continuous developmental transition, we took a scrape of a feeding front of NC4 cells, from inside the bacterial zone through to the mound stage of development. Cells were immediately inoculated into ice-cold KK2 buffer (20mM KPO_4_, pH 6.0), and disaggregated by gentle pipetting. To remove the bacteria, cells were centrifuged at 720g for 2 minutes, then resuspended in ice-cold KK2. Single cell transcriptomes were derived using a Chromium Single Cell A Chip platform (PN-1000009), with sequencing carried out using a NextSeq500 Mid-output 150-cycle kit. The protocols were as described previously (Nichols et al., 2020), with one modification, 12 and 13 cycles were used for sample index PCR of the replicates, rather than 11 and 12.

Alignment, barcode counting, UMI counting, and filtering was performed by Cell Ranger v3.1.0 using default parameters. For the two biological replicates, totals of 2925 and 2404 single cell libraries passed the filter, with a median of around 15000 and 13000 transcripts per cell (unique molecular counts: UMIs), and 2200 and 2300 genes per cell, for replicate one and two, respectively. All data analysis, unless otherwise stated, was performed in R. We excluded outliers with extremely high total UMI counts (1.5 interquartile ranges above the third quartile), cells with less than 3000 total UMI counts, and cells with less than 800 mapped genes. A total of 2671 and 2072 cells, from replicate one and two, respectively, were used for further analysis. Sequencing data are available at GEO: GSE220242

Molecular counts of cells were normalised using size factors calculated with ‘scran’ package (Lun et al., 2016). Dimensionality reduction was performed using genes with mean normalised UMI counts above 0.01 (9698 genes). Reducing the dimensionality of transcriptome data to two dimensions used principal component analysis (PCA) first, then elastic embedding on the first 11 principal components (selected based on their variance contribution), as described in (Chen et al., 2019). Elastic embedding was performed in MATLAB. For clarity, cells with higher expression are plotted on top of cells with lower expression. The set of aggregation-specific genes is the same as in (Nichols et al., 2020). The genes being up-regulated during the mound stage (“post aggregation”) and cell-cycle genes are as in (Antolovic et al., 2019). For all gene sets, we used an additional selection of genes having more than 10 captured transcripts in at least one cell in our dataset, except for cell-cycle genes having more than 1 transcript in at least one cell. The 3D cell density landscape was calculated with the use of ‘ks’ package in R, using the Hpi (bandwidth estimator) and kde functions. 2D cell density landscapes were plotted with the ‘ggplot2’ package, with the bandwidth set to 1. The cell-cell correlation matrix and two-way hierarchical clustering were carried in MATLAB, using the clustergram function.

To test the effects of loss of the adenylyl cyclase required for chemotaxis and aggregation (ACA) on the jump, we first identified the sets of genes being repressed at the jump (pre-jump), transiently expressed at the jump (jump), and upregulated after the jump (post-jump). We established criteria that generated consistent scRNAseq profiles for each category. The cells were divided into pre-jump, jump and post-jump based on the hierarchical clustering of the cell-cell correlation matrix (Fig. S4A). Jump class genes were defined as having fold change (FC) > 3 from before and after the jump, having the mean expression >1 normalised UMIs in the jump region and <10 before and after. Pre-jump genes were defined as being downregulated twofold in the jump region and tenfold after the jump (compared to the pre-jump region), with mean expression >10 normalised UMIs before and < 3 normalised UMIs after the jump. Post-jump genes were defined as having FC > 10 from before the jump and FC > 0.1 from the jump region, with mean expression < 5 UMIs before and >5 UMIs after the jump. For comparing the behaviour of these sets of genes in populations with or without ACA, we used the population RNAseq data from (Katoh-Kurasawa et al., 2021). For *aca^-^* expression values, we used the mean value of both replicates from this published dataset. For wild-type expression values we used mean value of replicates named r5, r6 and r7 (from the published data). We chose these replicates as they were sequenced on Illumina HiSEq 2500, the same as their ACA mutants. The genes we have considered here are those, in wild-type cells, that show the same temporal behaviour in our data (the wild isolate NC4 cells cultured on bacteria) and the earlier study (the lab strain Ax4, cultured in axenic media). A minor fraction of genes in each class (pre-jump 7/61, jump 8/30, post-jump 1/83) did not overlap between the datasets.

### Imaging protocols

Cells were harvested from adherent cultures in HL5 medium (Formedium), centrifuged at 720 g for 2 minutes and resuspended in 5 mL KK2. Cells were recentrifuged and resuspended at 1x10^7^ cells/mL in KK2. Transcriptional reporter and signalling reporter cells lines were mixed at a 1:2 ratio, and spotted onto an agar plate with diluted SM (1 SM : 19 H_2_O) freshly spread with *Klebsiella*. Agar plates were incubated upright in a humid chamber for 3 days when agar pads were excised and inverted onto a 35mm imaging dish (µ-Dish, Ibidi, 81156). Cells were imaged on an inverted spinning disk confocal microscope (3i) using a 63 x oil immersion lens, with a Prime 95B CMOS camera (Photometrics). 14 to 16 z slices were acquired with a 0.4 µm step size, with 2x2 binning. 3D stacks were captured every 45 or 60 seconds at multiple xy positions across the cell population. 9 fields of view were stitched together, using Slidebook (3i), to generate a montage of the entire heterogeneous early developmental niche. GFP and mCherry were excited with a 488 nm and 561 nm lasers, respectively, with laser powers optimised for best resolution alongside maintained cell health. Laser power, gain and exposure time were kept consistent between wildtype and *acaA*-cells of the same transcription reporter. Wildtype cells were imaged for 4 hours to capture the full range of cAMP signalling dynamics. The *acaA*-cells, which lack excitable cAMP signalling were imaged for 1.5 hours. For bPAC activation, transcriptional reporter cells were mixed with bPAC expressing cells at a 1:2 ratio. Activation used a 3D stack with a 488 nm laser every 6 minutes to mimic normal cAMP pulsing. This approach had the dual function of light activating bPAC and collecting transcription spot data. 12-15 fields of view were captured to sample the extended starvation zone of the *acaA-* cells.

### Spot tracking and image analysis

Spot detection was carried out as described (Corrigan et al., 2016) with the modifications that a threshold was imposed to distinguish between spots representing sites of transcription and background. Thresholds were established iteratively by inspection, then the same threshold was used for wild-type and *acaA-* experiments with that transcriptional reporter. Thresholds were applied to the background intensity of the nucleus, discarding very bright and very dim cells, which show non-linear scaling between live transcription site data and smFISH-based intensity measurements. To plot transcription dynamics in the colony through time and show the broadest dynamic range of the data, values above 3 inter-quartile ranges from the upper quartile were set to the highest colour. Pixels with no detected spots are set to black. To compare wild-type and *acaA-* cells, plots for wild-type and *acaA-* data were set to the same colour scale. To identify cAMP waves, we masked signal from the transcriptional reporter cells, which are more variable in their background intensity than the Flamindo2 cells. The intensity of the remaining cell-containing pixels (representing primarily the Flamindo2 signal) were averaged at each timepoint.

### Analysis of spatial signalling and transcription data

To measure jump gene expression, we analysed data from the population before the clear physical separation of the population into starving and streaming cells. The time of population separation was specified manually. Plots showing time series of nuclei counts and spot intensity were produced by plotting nuclear regression fits and spot intensity regression fits for each timepoint. To explore the relationship between cell position, cAMP signalling and jump gene transcription, images were divided into left and right regions, corresponding to recently starved cells and more starved cells. Using R, a local polynomial regression fit (loess function) was applied to nuclei counts across the x-dimension for each timepoint with a span of 0.75. All further local polynomial regression fits (ie. for Flamindo2, spot intensity) were carried out in the same way. The x positions of the minimum of the fit were identified for each timepoint. The range of values of the identified minima positions was used to define the least occupied area, i.e. the approximate location of cell population separation. The region spanning from the edge of the field occupied with bacteria to the edge of the least occupied area was defined as ‘left’ and the region spanning from the other edge of the least occupied area to the far right of the captured field as ‘right’. Mean values of transcription spot intensity and Flamindo2 signal were compared across the left and right regions of the image at each timepoint.

Inflections in the transcription spot intensity fit for each timepoint were used to characterise the location of cells where gene expression is strongly induced. To capture the dynamic location of the inflection over time, the x coordinate of the inflection was taken, and a local polynomial regression curve fitted linking inflection x coordinates through time, excluding outliers >1.5 interquartile ranges beyond the upper and lower quartiles.

Difference images of Flamindo2 were produced by taking the difference in intensity between one timepoint and the subsequent one, referred to as diff(Flamindo2). As the lowest intensity of Flamindo2 corresponds to the highest intracellular cAMP concentration, the timepoints where cAMP waves pass through the population were found by identifying local minima of mean d(Flamindo2) in the right-hand region of the image at each timepoint. To find the border of cAMP relay, a local polynomial regression fit was applied to diff(Flamindo2) in x, and inflections in the fit were found for timepoints previously identified as having a cAMP wave. To capture the location of the cAMP signal relay border in time, the x coordinate of the inflection was taken, and a local polynomial regression curve fitted, linking inflection x coordinates through time excluding outliers >1.5 interquartile ranges beyond the upper and lower quartiles.

To investigate the time delays between signalling and transcription, transcription spot intensity mean and Flamindo2 mean signal in the right region of the image were further processed. Using R, both signals initially were locally smoothed using Savitzky–Golay filtering, with a filter length of 3 and a filter order of 1. We then corrected for the changing background over time, by Savitzky–Golay filtering with a filter length of 15 and a filter order of 1. The background was then subtracted from the locally smoothed signal. Flamindo2 signal is inversely related to [cAMP], so to convert Flamindo2 signal into cAMP signal Flamindo2 intensity was inverted. To find the time difference between cAMP and gene transcription dynamics, cross correlation analysis was completed on the smoothed, background corrected signals. Cross correlation was completed over a period of +/-10 minutes to find realistic signal relay times without artefacts from noisy signal.

## Supplementary Information

### Supplementary Movies

**Movie 1**. Imaging jump transcription over the length scale of the developmental niche. The time series progressively zooms out, showing transcriptional events in single cells, then zooming out to show the position and movement of individual cells in the context of the early developmental niche. The cells are a mixture of *cafA* transcriptional reporter cells and cells expressing the cAMP reporter, Flamindo2. The contrast has been optimised in this case to make the transcriptional reporter cells clearly visible. In the first frame, the arrow points to a cell with a bright nuclear spot corresponding to nascent *cafA* RNA, at the site of transcription. As the movie zooms out, the full niche becomes apparent, with the undifferentiated cells to the left and the differentiating cells further to the right, with the onset of cell aggregation (streaming) visible. The playback is at 7 frames per second, with each frame captured every 1 minute. When fully zoomed out, the field of view is 1.68mm along the long axis.

**Movie 2**. Imaging cAMP signalling dynamics in the developmental niche. This move shows similar data to Movie 1, with the contrast optimised to show the dimmer Flamindo2 expressing cells. The undifferentiated, feeding cells are on the left, with the cell aggregation beginning on the right. Towards the right, the population shows synchronous oscillations in Flamindo2 intensity, corresponding to cAMP oscillations (Flamindo2 signal is inversely related to intracellular cAMP level). The playback is at 15 frames per second, with each frame captured every 45 seconds. The field of view is 1.68mm along the long axis.

---

**Supplementary Figure 1.**
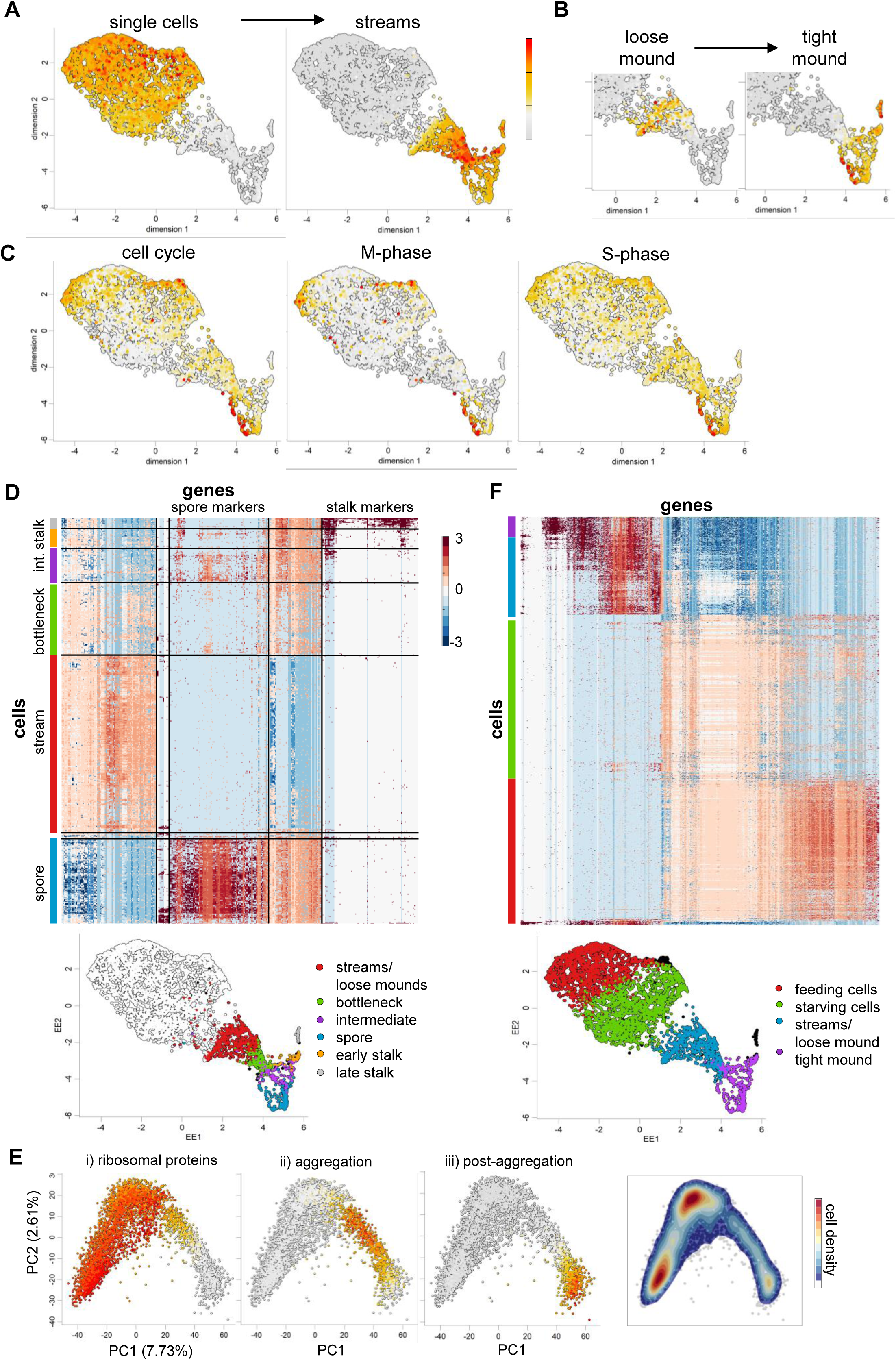
Developmental progression in the physiological niche. **A** and **B)** Cross referencing the 2D scRNAseq cell map to previously published population RNAseq data: genes identified as markers of major morphological changes in (Katoh-Kurasawa et al., 2021). In A, the genes changing during the transition from single cells to cell streams (down-regulated: 140 genes, up-regulated: 91). In B, the genes changing during the loose to tight mound transition (down-regulated: 4, up-regulated: 83). **C)** Distribution of cell-cycle genes expression on the 2D map (160 genes). For optimal visual contrast, the maximal colour value here is set at mean expression > 0.2. Plots are also shown for M-phase and S-phase genes. The latter shown more diffuse enrichment. **D)** Two-way hierarchical clustering of 1112 cells from the post-jump region (selected as the cells belonging to the right main branch in Fig.S4A). Colour shows z-score values. Genes were selected as having mean normalized expression over 0.01 and being correlated with at least ten other genes with Pearson’s |r| > 0.5. Gene clusters expressing spore and stalk markers are indicated at the top of the map. Six main clusters of cells are marked with coloured boxes left of the heatmap. These six clusters are overlaid on the adjacent 2D transcriptome plot. The clusters are streams/loose mounds, bottleneck, intermediate, spore and two stalk populations. **E)** PCA plots labelled with the different stage gene sets used in Figure 1B, showing that the PC1 axis records developmental time. **F)** Two-way hierarchical clustering of cells described with 957 genes (colour shows z-score values). Genes were selected as in D. Four main clusters of cells are marked with coloured boxes left of the heatmap. These four clusters identified are overlaid on the 2D transcriptome plot. The main clusters are feeding cells (1624), starving cells (1840), streams/loose aggregate (737) and tight aggregate (396). 66% of the identified genes are repressed before the jump.

**Supplementary Figure 2.**
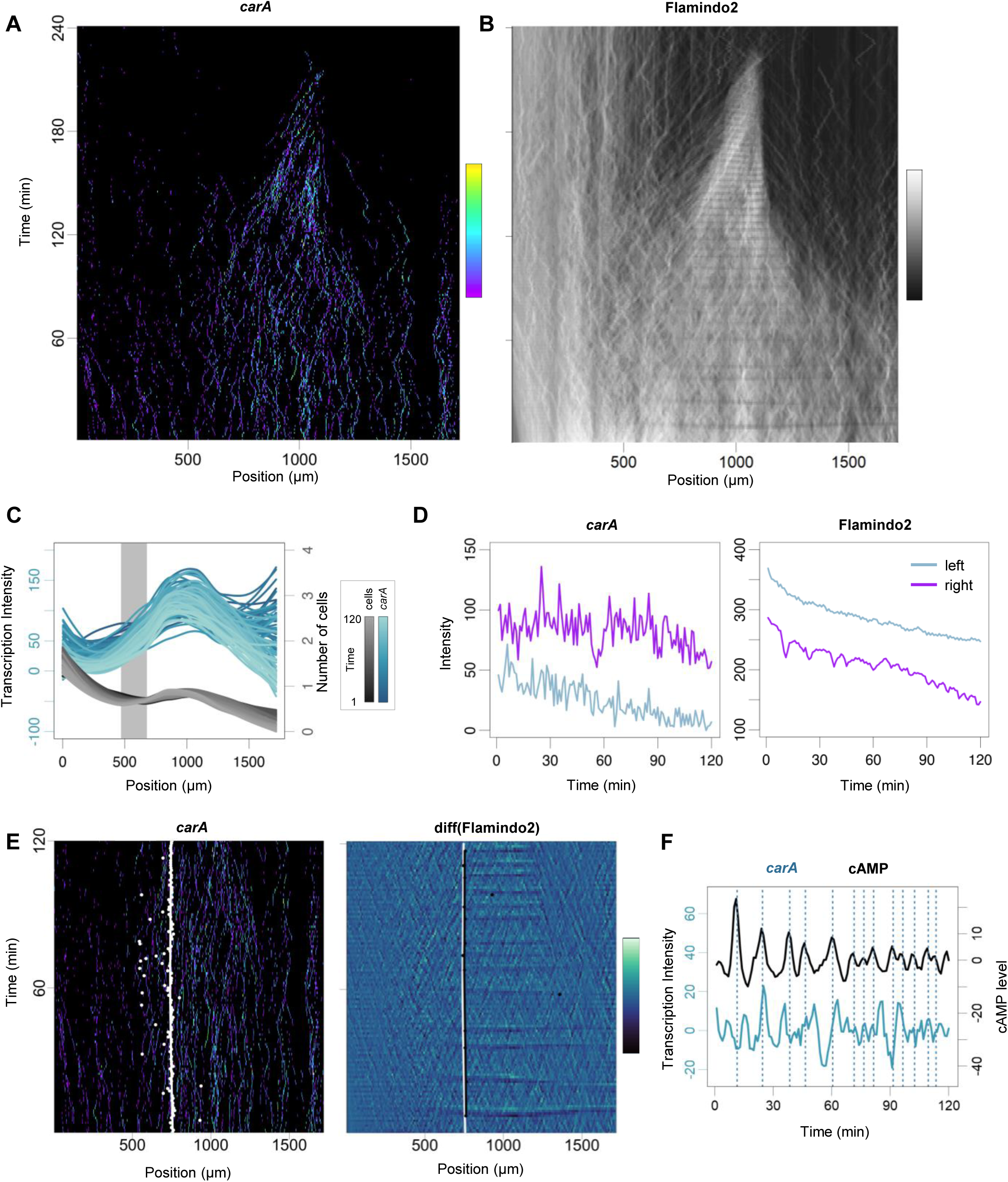
Coupling between jump transcription and signalling dynamics. This figure shows a repeat experiment of Figure 2. **A)** Imaging transcription dynamics of the jump gene, *carA*. Horizontal axis reflects the axis of differentiation. Imaging time is represented on the vertical axis. Transcription activity (detected using PP7-PCP) over time is shown, with activity level related by the colour scale on the right. Transcription is initially rare and sporadic, becoming frequent and oscillatory as differentiation proceeds. **B)** Same data as in A, showing cAMP signalling dynamics using the Flamindo2 biosensor, which dims in fluorescence when intracellular [cAMP] increases. Data show oscillations in differentiating cells. Cells merge into an aggregate during the movie. **C)** Increased transcription activity during differentiation. Plots summarises data in B, and also shows the distribution of cells in the population. Changing transcription and cell distributions over time are shown as different colour shades (see colour scale). The grey line corresponds to the minimum in cell density, which corresponds to where the population splits during the transition to multicellularity. **D)** Transitions in transcription and signalling dynamics. Left panel shows the differences in *carA* transcription dynamics between the areas left and right of the grey line in D. Right panel shows the differences between oscillatory and non-oscillatory cAMP dynamics either side of the grey line in D. **E)** Positional coupling between transcription and signalling dynamics. Left-white spots represent inflections of the curves of transcriptional intensity values at each imaging time point. The white line is a regression line summarising the distribution of points. Right-black dots show inflection points for cAMP signalling, where the signal intensity changes at the left extremity of the waves. Black line is the regression summarising these points. White line the same as in left panel. Inflection values were calculated at timepoints corresponding to cAMP wave maxima. **F)** Temporal coupling between transcription and signalling oscillations. Peaks in cAMP signalling (vertical lines) 4-5 minutes prior to peaks in *carA* transcription.

**Supplementary Figure 3.**
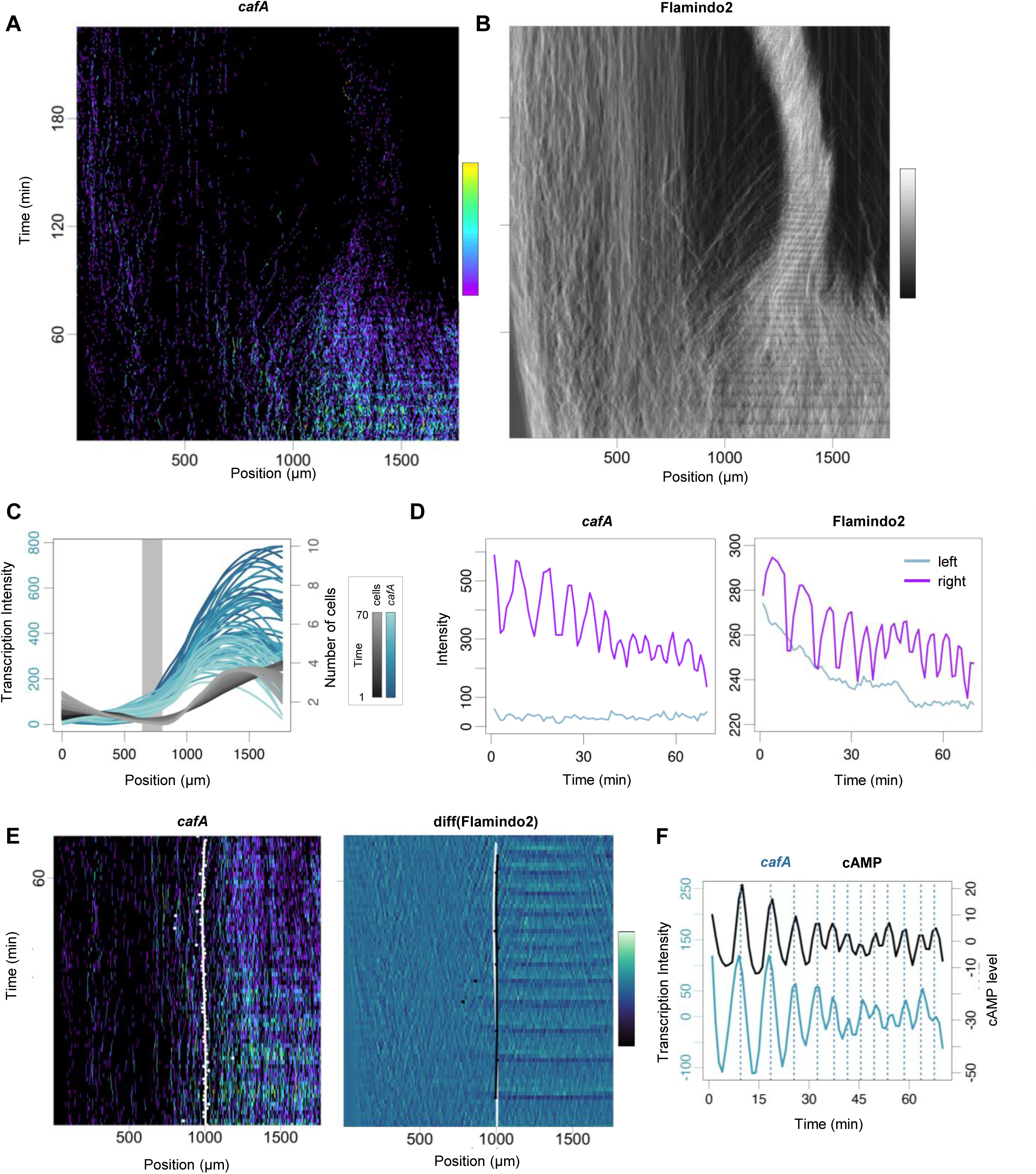
Alternative coupling strategies between jump signalling and transcription. This figure shows a repeat experiment of Figure 3. **A)** Imaging transcription dynamics of the jump gene, *cafA*. Horizontal axis reflects the axis of differentiation. Imaging time is represented on the vertical axis. Transcription activity measured using the PP7-PCP system. Transcription is initially rare and sporadic, becoming frequent and oscillatory as differentiation proceeds. **B)** Same data as in A, showing cAMP signalling dynamics using Flamindo2. Cells merge into an aggregate during the movie. **C)** Increased transcription activity during differentiation. Plots summarises data in A, and shows the distribution of cells in the population. Changing transcription and cell distributions over time are shown as different colour shades (see colour scale). The grey line corresponds to the minimum in cell density, which corresponds to where the population splits during the transition to multicellularity. **D)** Transitions in transcription and signalling dynamics. Left panel shows the distinction in *cafA* transcription dynamics between the areas left and right of the grey line in C. Right panel shows the distinction between oscillatory and non-oscillatory cAMP dynamics either side of the grey line in C. **E)** Boundaries of transcription and signalling dynamics. Left-white spots represent inflections of the curves of transcriptional intensity values at each imaging time point. The white line is a regression line summarising the distribution of points. Right-black dots show inflection points for cAMP, where the signal intensity changes at the left extremity of the waves. Black line is the regression summarising these points. White line the same as in left panel. Inflection values were calculated at timepoints corresponding to cAMP wave maxima. **F)** Temporal coupling between transcription and signalling oscillations. Peaks in cAMP signalling slightly precede peaks in *cafA* transcriptional activity.

**Supplementary Figure 4.**
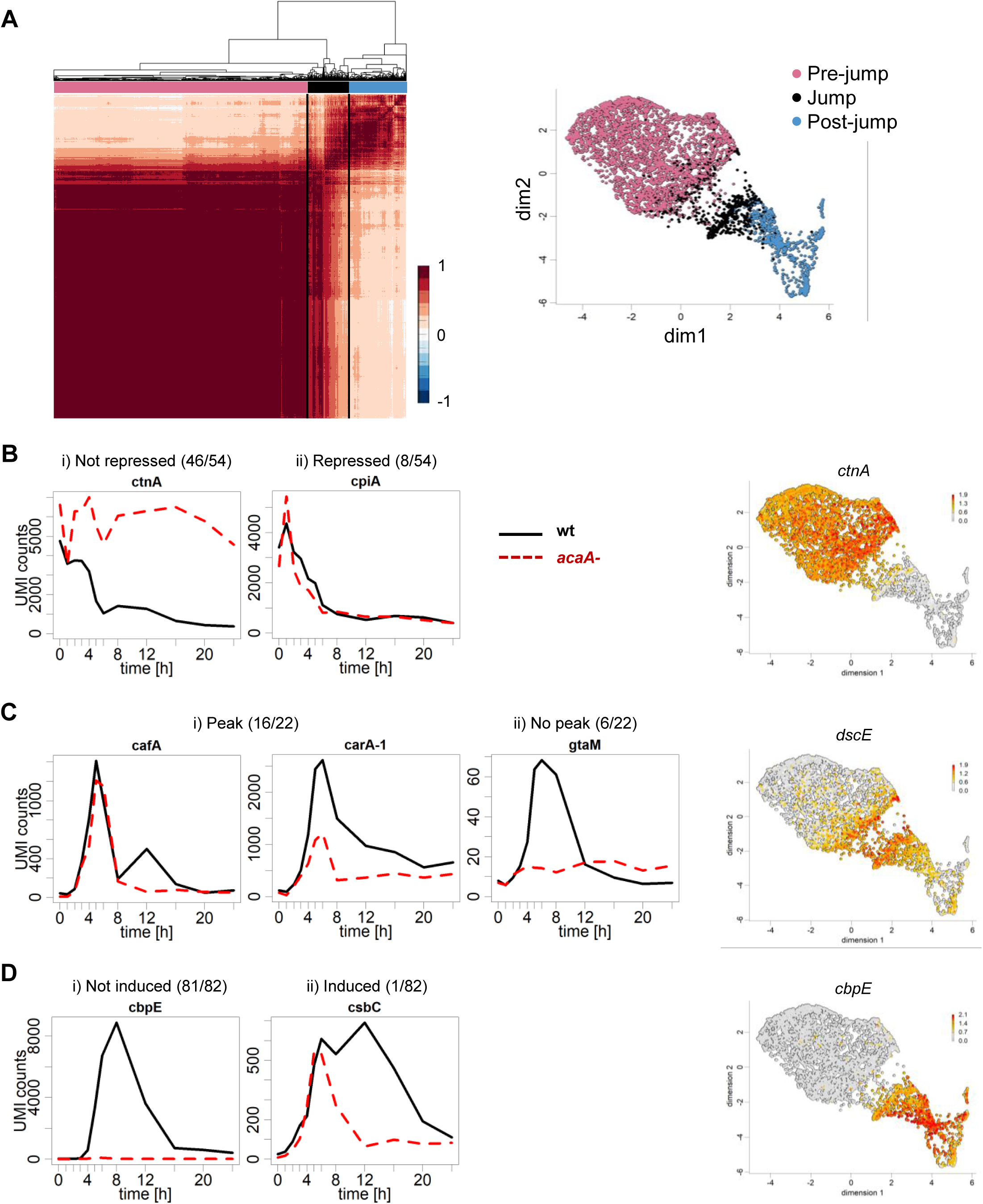
Changes in jump gene expression caused by loss of cAMP. **A)** Cell-cell correlation matrix of the continuous developmental scRNAseq (the same plot as Fig. 1D, but including the hierarchical tree). We identified three main clusters, with the one in the centre corresponding to the jump. To find jump genes, we expanded this region by taking the most-right subcluster of the left cluster and the most-left subcluster of the right cluster (black box). The modified clusters (before, during and after the jump) marked as (pink, black and blue) are shown in the right panel on the 2D transcriptome plot. We used this division to identify genes downregulated before the jump (**B**), enriched in the jump region (**C**) or upregulated after the jump (**D**). **B – D** show transcript dynamics of genes identified as pre-, during and post-jump, in wild-type cells (black line) and *aca^-^* cells (red dashed line), which lack the aggregation stage cAMP synthesis enzyme. Transcript data from (Katoh-Kurasawa et al., 2021). The proportion of the genes showing each specific behaviour are shown. Also shown are typical examples of scRNAseq expression plots for each class. **B)** i) Most transcripts normally downregulated at the jump retain expression in *acaA-* mutants (85%). ii) A minor proportion of jump repressed transcripts are unperturbed. **C)** i) 73% of jump-specific transcripts retain a peak in expression in *acaA-* mutants, although induction is reduced, ii) 27% of jump specific genes are not induced in *acaA-* cells. **D)** Almost all (99%) genes induced after the jump are not induced in *aca^-^* mutants.

## Notes

### Competing Interest Statement

The authors have declared no competing interest.

